# Sphingosine induction of the *Pseudomonas aeruginosa* hemolytic phospholipase C/sphingomyelinase, PlcH

**DOI:** 10.1101/2023.11.06.565848

**Authors:** Jacob R. Mackinder, Lauren A. Hinkel, Kristin Schutz, Korin Eckstrom, Kira Fisher, Matthew J. Wargo

**Affiliations:** Department of Microbiology and Molecular Genetics, Larner College of Medicine, University of Vermont; Cellular, Molecular, and Biomedical Sciences Graduate Program, University of Vermont

**Keywords:** sphingolipid, lipid, pathogenesis, virulence, transcription regulation

## Abstract

The hemolytic phospholipase C, PlcH, is an important virulence factor for *Pseudomonas aeruginosa*. PlcH preferentially hydrolyzes sphingomyelin and phosphatidylcholine and this hydrolysis activity can drives tissue damage, inflammation, and interferes with the oxidative burst of immune cells. Among other contributors, transcription of *plcH* was previously shown to be induced by phosphate starvation via PhoB and by the choline metabolite, glycine betaine, via GbdR. Here, we show that sphingosine can induce *plcH* transcription and resultant secreted PlcH enzyme activity. This induction is dependent on the sphingosine-sensing transcription regulator SphR. The SphR induction of *plcH* occurs from the promoter for the gene upstream of *plcH* that encodes the neutral ceramidase, CerN, and transcriptional read-through of the *cerN* transcription terminator. Evidence for these conclusions come from mutation of the SphR binding site in the *cerN* promoter, mutation of the *cerN* terminator, enhancement of *cerN* termination by adding the *rrnB* terminator, and RT-PCR showing that the intergenic region between *cerN* and *plcH* is made as RNA during sphingosine, but not choline, induction. We also observe that, like glycine betaine induction, sphingosine induction of *plcH* is under catabolite repression control, which likely explains why such induction was not seen in other studies using sphingosine in rich media. The addition of sphingosine as a novel inducer for PlcH points to regulation of *plcH* transcription as a site for integration of multiple host-derived signals.

## Introduction

The secreted hemolytic phospholipase C/sphingomyelinase, PlcH, was one of the earliest identified *Pseudomonas aeruginosa* virulence factors^1^. PlcH preferentially hydrolyzes phosphatidylcholine and sphingomyelin to yield phosphorylcholine and either diacylglycerol or ceramide, respectively^2^. Unlike virulence systems commonly lost during chronic infection^3-5^, PlcH production is retained in all clinical and environmental strains that have been reported, though levels vary between strains^2,6-8^. Experimentally, *plcH* mutants have decreased virulence in many infection models and strains producing higher levels of PlcH cause more lung damage^8-12^, due to a combination of cytotoxicity, pro-inflammatory properties, and surfactant damage^9,13-17^. As befits a secreted enzyme that likely functions as a public good, *plcH* transposon insertants do not typically have substantial competition defects in transposon sequencing studies^18-21^, as mutants are presumably rescued by community production of PlcH. In humans, PlcH production is negatively correlated with patient outcomes and treatment efficacy in CF^6,7^.

Like many important virulence pathways, there are a number of inputs into PlcH regulation. GbdR activates transcription in response to glycine betaine (GB) or dimethylglycine and binds directly to the *plcH* upstream region ∼90 bases upstream of the GB-dependent transcription start site^22-27^. GB and dimethylglycine can lead to *plcH* induction directly, whereas choline can only induce *plcH* when metabolized to GB via the BetBA enzymes^22,27^. Transcription of *plcH* is also induced under conditions of phosphate starvation, transduced through the PhoBR system^25^. Additionally, Anr can function as a repressor of *plcH* transcription under varying oxygen conditions^28^ in response to metabolites from the GB catabolic pathway^29^. Induction of *plcH* by GB is under catabolite repression control^26^, but it is not known if that is direct or indirect. There is also a Vfr binding site predicted ∼300 bases upstream of the *plcH* translational start site, however, no function arising from this site has been shown and Vfr was previously ruled out as the mediator of *plcH* catabolite repression^26^. The H-NS like proteins MvaU and MvaT are known to directly or indirectly regulate expression from many loci in Pa, including *plcH* ^30^. PhrS has also been shown to negatively regulate *plcH* expression, likely indirectly^31^.

In this study, we add to the already multifaceted transcriptional regulation of *plcH* by showing that the host-derived sphingolipid, sphingosine, is a novel inducer of *plcH* transcription and resultant extracellular PlcH. Rather than acting directly on the *plcH* promoter like GbdR and PhoB, sphingosine induction is the result of activation of the *cerN* promoter via SphR and transcriptional read-through from *cerN* to *plcHR*. This regulation was not previously observed when sphingosine induction of *plcH* was examined^32^ because, as we show here, sphingosine induction of *plcH* is under catabolite repression control. Together with previously described regulatory pathways, these findings highlight *plcH* as a node for integration of important host-derived metabolites.

## Results

### Sphingosine promotes PlcH production via transcriptional induction

In unrelated studies to understand sphingosine-dependent transcriptional responses in multiple bacterial species, we conducted sphingosine induction of *P. aeruginosa* PAO1 in minimal media and observerd that reads from *plcH* and *plcR* were much more abundant in the presence of sphingosine. The *plcHR* operon encodes the hemolytic phospholipase C/sphingomyelinase, PlcH, and its chaperone, PlcR. The *plcHR* operon is transcriptionally downstream of the *cerN* gene, which encodes the neutral ceramidase^33,34^ (**Fig. 1A**). As sphingosine was not a previously known inducer of PlcH, we chose to further examine this induction route.

**Figure 1:**
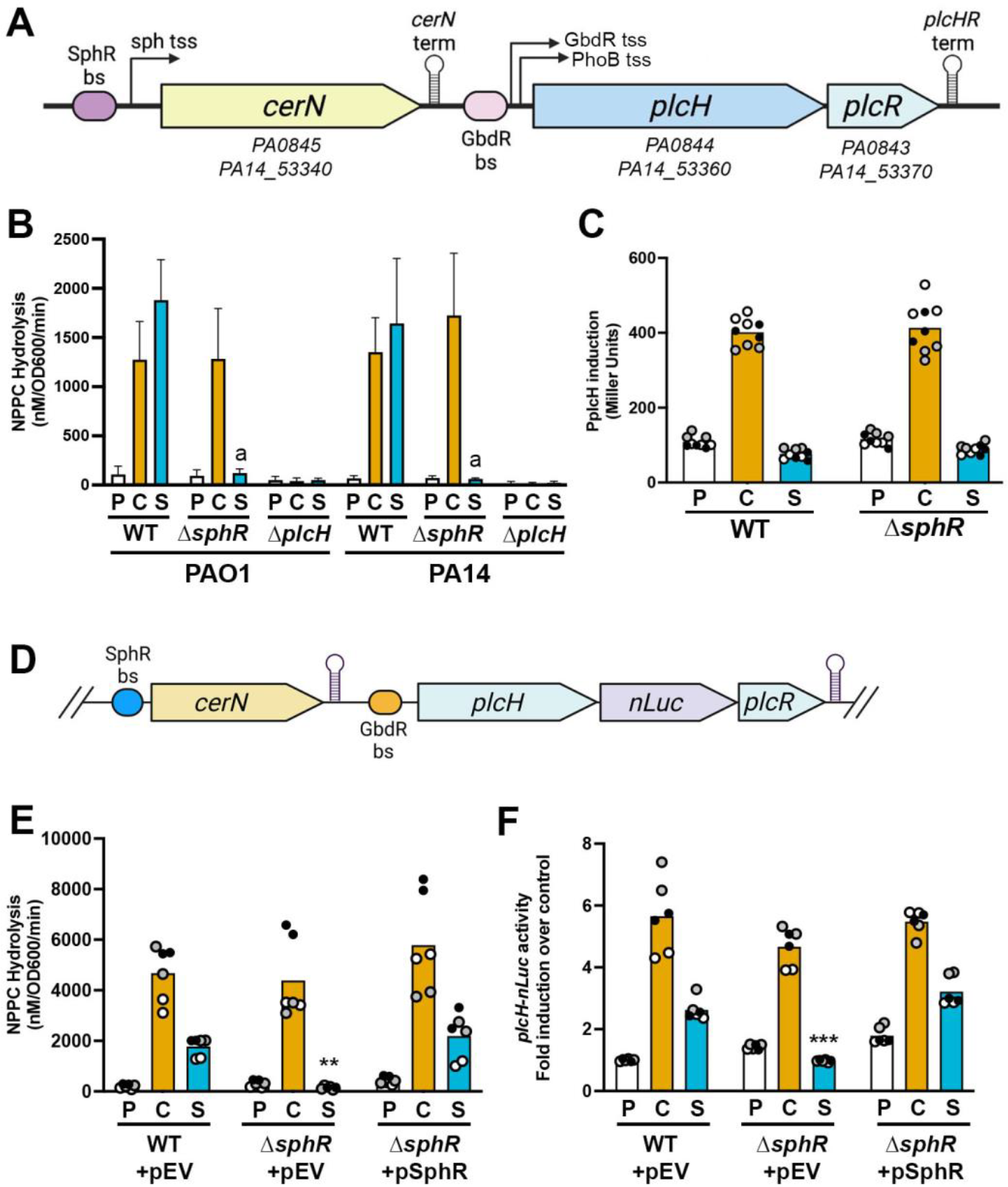
Spingosine induction of PlcH. **(A)** Genomic region containing *plcH, plcR*, and *cerN* with gene numbers from both PAO1 and PA14 with components important for this study diagrammed and labeled. **(B)** Sphingosine induces extracellular PlcH activity in both PAO1 and PA14 as measured by nitrophenylphosphorylcholine (NPPC) hydrolysis. The “a” denotes p < 0.0001 comparing Δ*sphR* to WT in the presence of sphingosine using 2-way ANOVA with Tukey’s post-test. All Δ*plcH* conditions are significantly lower than WT and Δ*sphR* in both strains. Error bars represent standard deviation **(C)** β-galactosidase activity from a P_*plcH*_-*lacZYA* plasmid-encoded reporter (pMW22) showing that choline leads to induction in both WT and Δ*sphR*, while sphingosine does not alter expression and loss of *sphR* has no impact (2-way ANOVA with Sidak’s post-test). **(D)** Diagram of the chromosomal locus for strains examined in Figs 1E and 1F showing the integration of the nano-luciferase open reading frame in between *plcH* and *plcR*, generating a synthetic *plcH-nLuc-plcR* operon. **(E & F)** Sphingosine induction of PlcH activity and transcription requires *sphR* and loss of *sphR* on the chromosome can be complemented by *sphR* expressed from a plasmid (pSphR) for both PlcH enzyme activity (E) and nLuc activity (F). Statistical significance noted as ** (p < 0.01) and *** (p < 0.001) using a 2-way ANOVA with Tukey’s post-test and the noted comparisons are to the WT pEV in the respective condition. The *sphR* complement is not significantly different than WT in any condition. For panels C,E,F, all data points are shown and are colored by experiment with white circles for all replicates from experiment #1, grey from experiment #2, and black from experiment #3. Only the means for each experiment are used in the statistical analyses for these panels (n = 3 per condition). Gene organization diagram generated with BioRender. Abbreviations: bs, binding site; sph, sphingosine; tss, transcription start site; term, transcriptional terminator; P, pyruvate (control); C, choline; S, sphingosine; pEV, empty vector pMQ80.

We first tested if sphingosine induction of *plcHR* lead to increased secreted PlcH as measured by enzymatic assay based on nitrophenylphosphorylcholine (NPPC) hydrolysis. **Figure 1B** shows that choline and sphingosine can both independently drive increased PlcH enzyme activity in the supernatant and that sphingosine induction of PlcH activity was not observed in an *sphR* deletion. We had previously used the region immediately upstream of the *plcH* gene as a negative control for EMSAs testing SphR binding^35^ and, in support of that, the P_*plcH*_*-lacZYA* transcriptional reporter shows no response to sphingosine but responds to choline induction (**Fig 1C**), as we have previously shown^23^. We then integrated a codon-optimized nanoluciferase gene (*nLuc*) into the *plcHR* locus to function as a chromosomal reporter: *plcH-nLuc-plcR* (**Fig. 1D**). Integration of nLuc at this position did not alter PlcH secreted activity to either sphingosine or choline (**Fig. 1E** and Supplemental DataSet2) and measurement of the chromosomally-encoded *nLuc* showed induction by sphingosine (**Fig. 1F**). Though the pattern of nLuc expression mirrors PlcH activity, nLuc appears to be very stable and accumulates in cells during overnight growth, which decreases the dynamic range of this reporter when starting from the high cell inoculum we use to simultaneously measure supernatant NPPC hydrolysis activity within this short timeframe.

### Sphingosine induction of *plcH* transcription is due to transcriptional readthrough from the P_cerN_ promoter

Upstream of the *plcHR* operon is *cerN*, encoding a neutral ceramidase, which is a known target for sphingosine-dependent transcriptional induction^32^ controlled by SphR (**Fig. 1A**)^35^. To determine whether the overall organization of this region, including the *cerN* promoter, was needed for *plcH* induction by sphingosine, we tested a plasmid-borne P_*cerN*_-*fLuc*-P_*plcH*_-*nLuc* reporter (**Fig. 2A**, bottom). Wild-type cells carrying this reporter showed induction nano-luciferase activity in the presence of sphingosine was not induced in response to sphingosine in an *sphR* deletion mutant. Choline induction of nano-luciferase activity was not impacted by *sphR* deletion (data not shown).

**Figure 2:**
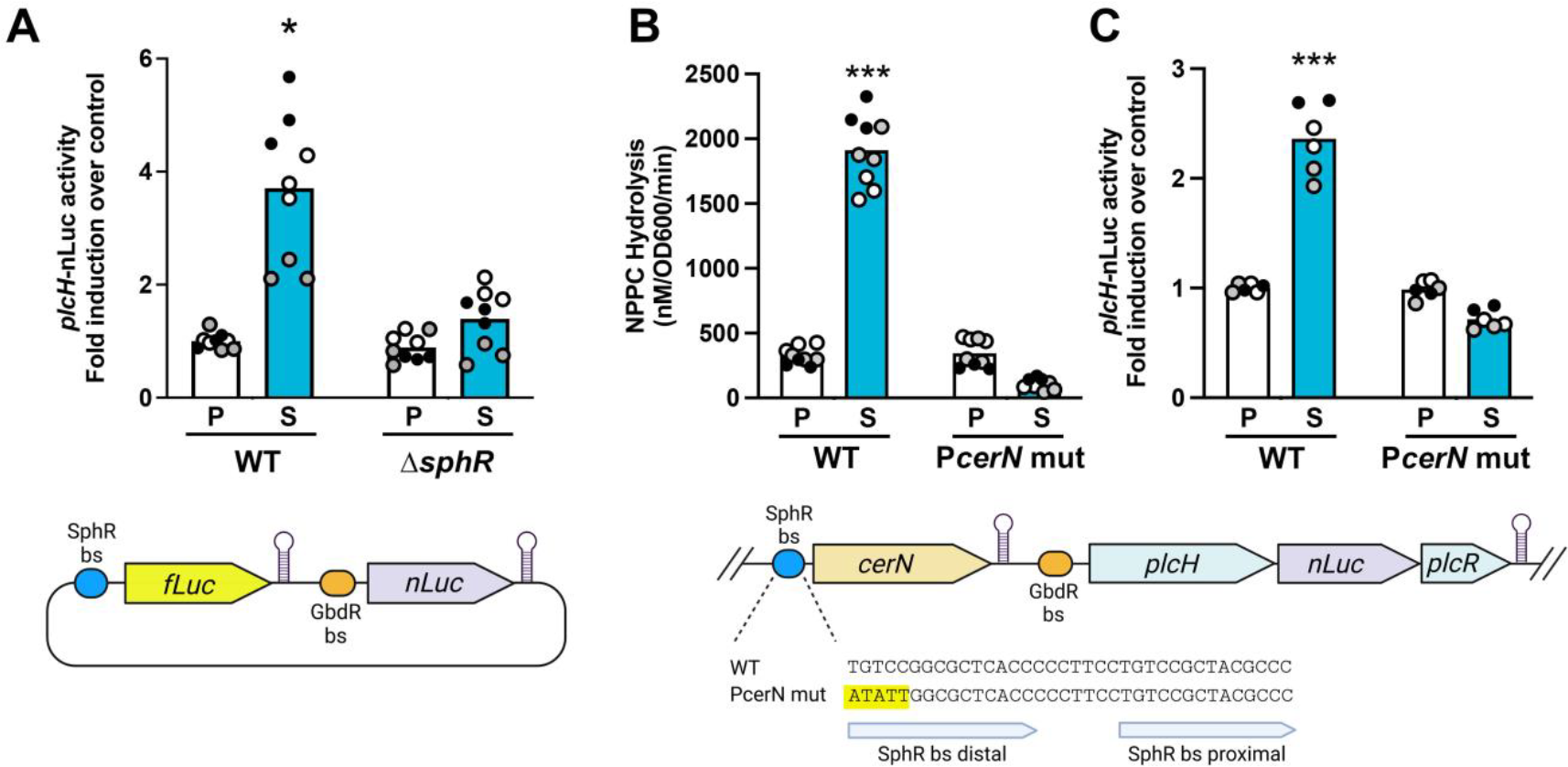
Sphingosine induction of PlcH/*plcH* dependent on the *cerN* promoter. **(A)** Dual luciferase assay results from strains carrying the plasmid diagramed below the graph. The plasmid results in a P_*cerN*_-*fLuc*-P_*plcH*_-*nLuc* dual reporter that mimics the chromosomal locus without the *cerN* or *plcH* coding sequences. Sphingosine induction from the *plcH* promoter is dependent on *sphR* with significance tested using 2-way ANOVA with Sidak’s post-test with * denoting p < 0.05. **(B & C)** Mutation of five bases in the distal SphR binding site in the chromosomal *cerN* promoter region (yellow highlighted residues) leads to loss of extracellular PlcH activity **(B)** and resultant *plcH-nLuc* activity **(C)**. Significance tested using 2-way ANOVA and Dunnett’s post-test with WT pyruvate as the comparator, with *** denoting p < 0.001. For all panels, all collected data points are shown and are colored by experiment with white circles for all replicates from experiment #1, grey from experiment #2, and black from experiment #3. Only the means for each experiment are used in the statistical analyses for these panels (n = 3 per condition). Gene organization diagram generated with BioRender. Abbreviations: P, pyruvate (control); C, choline; S, sphingosine; bs, binding site; mut, mutant.

Mutation of the SphR binding site in the *cerN* promoter at the native locus leads to elimination of sphingosine-dependent induction of extracellular PlcH activity and *plcH* transcriptional induction (**Figs. 2B-C**). Since sphingosine induction of *plcH* requires SphR regulation from the *cerN* promoter (**Figs. 2B-C**) and does not require any part of the *cerN* coding sequence (**Fig. 2A**) that strongly suggests the existence of a conditional operon the includes *cerN, plcH*, and *plcR* leading to a *cerNplcHplcR* polycistron.

To determine potential for transcriptional readthrough from the *cerN* promoter, we conducted reverse transcriptase PCR to examine the existence of RNA containing *cerN, plcH*, and the *cerN-plcH* intergenic region starting from the 3’ end of *cerN* open reading frame and continuing into the 5’ end of the *plcH* open reading frame (**Fig. 3A**). The RT-PCR supports the existence of this transcript and suggests that the *cerN* transcriptional terminator may not be a complete block to transcriptional extension.

**Figure 3:**
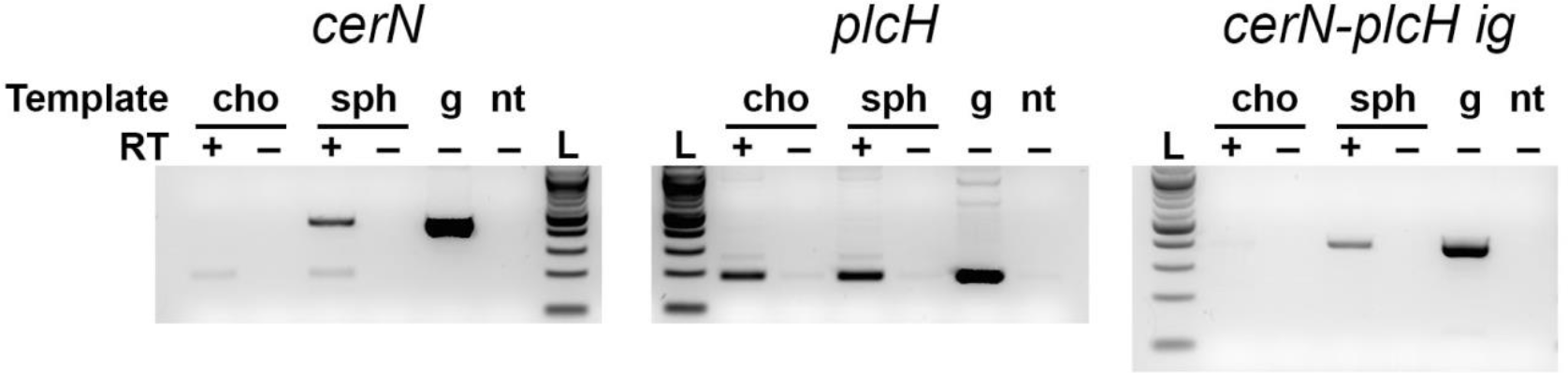
Evidence for conditional operon containing *cerN* and *plcH*. Reverse Transcriptase PCR demonstrates a contiguous RNA from the 3’ end of the *cerN* coding sequence to the 5’ end of the *plcH* coding sequence. The templates were total RNA isolated from choline induction (cho) or sphingosine induction (sph) with a positive control of genomic DNA (g) and a negative control without RNA. The inclusion of reverse transcriptase into the cDNA synthesis mix is noted in the RT label with + designating inclusion of reverse transcriptase. In agreement with all presented data and previously published work, the *cerN* coding sequence is detected only in RNA from sphingosine induction, the *plcH* coding sequence is detected in RNA from both choline and sphingosine induction, and the region spanning the *cerN-plcH* intergenic region (*cerN-plcH ig*) is detected only in RNA from sphingosine induction.

To test whether the predicted *cerN* terminator functioned as such, we tested its ability to prevent transcriptional readthrough using a synthetic operon on a plasmid containing P_*BAD*_-*gfpmut3*-ig_*cerNplcH*_-*nLuc* compared to a version that lacked the distal half of the predicted terminator stem loop. Mutation of the *cerN* terminator allowed completely permissive transcriptional read-through, while the wild-type *cerN* terminator greatly inhibited read-through (**Fig. 4A**). The presence of sphingosine did not impact the degree of termination (**Fig. 4A**) nor did heterologous testing in *E. coli* (data not shown) and thus the terminator appears to be unregulated in this context. To examine the impact of transcriptional terminator strength on sphingosine-dependent *plcH* induction, we inserted the *rrnB* transcriptional terminator immediately upstream of the native *cerN* transcriptional terminator on the chromosome. Adding this additional transcription terminator substantially reduced sphingosine induction of PlcH enzyme activity (**Fig. 4B**) and abrogated nano-luciferase induction from the *plcH-nLuc-plcR* locus (**Fig. 4C**). Together, these data support the function of the *cerN* terminator in the *cerN-plcH* intergenic region but suggest it likely functions as a static dose control system.

**Figure 4:**
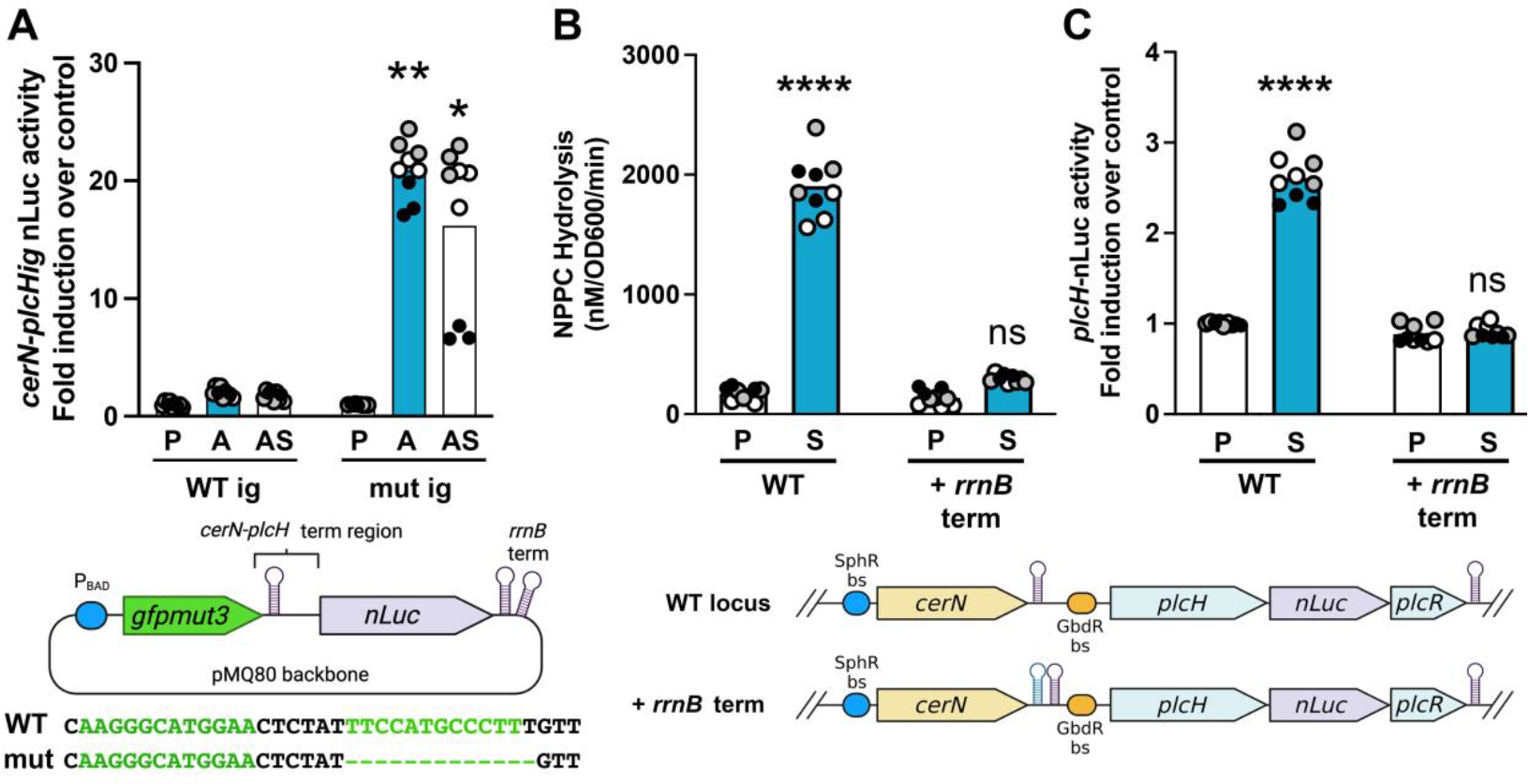
Role of the predicted *cerN* terminator in transcriptional termination. **(A)** The predicted *cerN* terminator does promote transcriptional termination. Expression of nLuc from a synthetic operon containing *gfpmut3* and *nLuc* with the wild-type *cerN* terminator or a mutant with the distal leg of the terminator stem loop deleted (see diagram below data). WT PA14 containing these plasmids were grown in a MOPS Pyruvate (P), with 0.1% L-arabinose (A), or with both L-arabinose and 100 μM sphingosine (AS). **(B & C)** Addition of the *rrnB* terminator upstream of the predicted *cerN* terminator leads to loss of significant extracellular PlcH activity **(B)** and resultant *plcH-nLuc* activity **(C)**. Significance tested using 2-way ANOVA and Dunnett’s post-test with WT pyruvate as the comparator, with *** denoting p < 0.001. For all panels, all of the collected data points are shown and are colored by experiment with white circles for all replicates from experiment #1, grey from experiment #2, and black from experiment #3. Only the means for each experiment are used in the statistical analyses for these panels (n = 3 per condition). Gene organization diagram generated with BioRender. Abbreviations: P, pyruvate (control); A, L-arabinose; S, sphingosine; bs, binding site; mut, mutant; ig, intergenic; term, terminator.

### Sphingosine induction of PlcH is under catabolite repression control

PlcH was shown to be unresponsive to sphingosine in a previous study conducted in a peptone and yeast-extract containing medium^32^, while our study uses a minimal medium, which made us suspect catabolite repression control. PlcH induction by choline is under catabolite repression control while PlcH induction by phosphate starvation is not^26^. To test if sphingosine:SphR dependent induction of PlcH production is under catabolite repression control, we measured NPPC production during growth in minimal media with pyruvate, glucose, or succinate as primary carbon sources. Both choline and sphingosine induction of PlcH activity are repressed during growth on glucose and succinate compared to pyruvate (**Fig. 5**). The responsible mechanism for this regulation is currently unknown but we speculate on possible mechanisms in the discussion.

**Figure 5:**
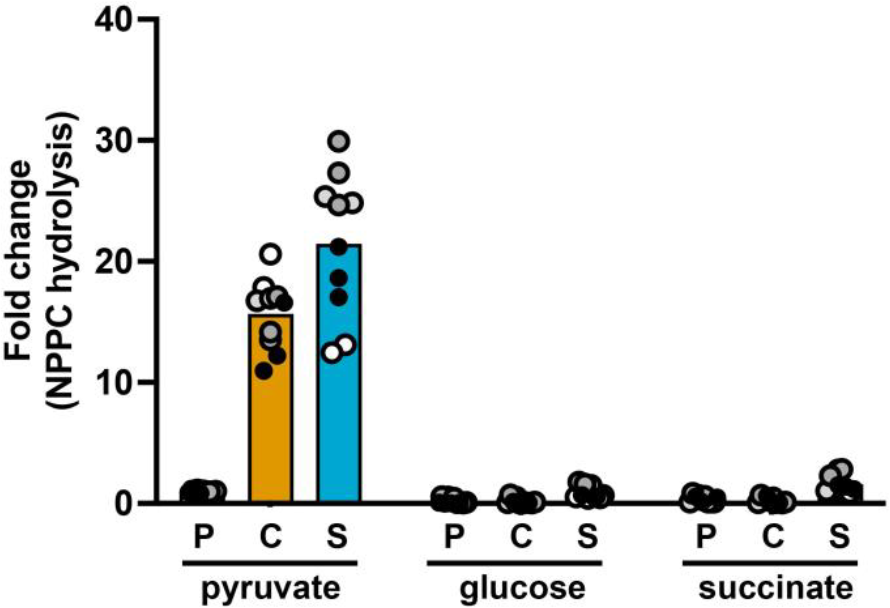
Extracellular PlcH activity is under catabolite repression control. Both choline and sphingosine induction of PlcH activity is repressed when cells are grown with either glucose or succinate as sole carbon sources. Pyruvate is permissive to PlcH induction by both compounds and is known to not trigger catabolite repression control. Analysis by 2-way ANOVA with Dunnett’s post-test using pyruvate induction in pyruvate media as the comparator (far left bar) shows that only choline and sphingosine induction in pyruvate media are significantly different (p < 0.0001). The trend toward induction for sphingosine in glucose and succinate media are small (<50% increase) and not statistically significant. All of the collected data points are shown and are colored by experiment, with white circles for all replicates from experiment #1, light grey from experiment #2, dark grey from experiment #3, and black from experiment #4. Only the means for each experiment are used in the statistical analyses for these panels (n = 4 per condition).

## Discussion

PlcH is an important virulence factor for *P. aeruginosa* capable of initiating inflammation, damaging pulmonary surfactant, and direct cell lysis^2,6,8,11,12,16,36-38^. Here, we identified an additional host-derived molecule driving PlcH induction – sphingosine. Sphingosine induction of *plcH* transcription is mediated by the sphingosine-binding regulator SphR in the context of transcriptional readthrough from the upstream *cerN* promoter. This was surprising given the previous report that sphingosine was unable to induce PlcH activity^32^. However, this is likely due to catabolite repression of PlcH expression in rich media as discussed further below. Sphingosine induction of PlcH is a second positive feedback loop controlling its expression, along with the GB-GbdR controlled positive feedback loop. Given the potential that these two systems, Anr regulation, and PhoB regulation may be functional at the same time in the host, an important ongoing question is how these signals interplay, integrate, or impact each other for ultimate production of PlcH during infection. Response to these signals could potentially be synergistic, interfering, temporally separated within a single cell, or physically separated amongst sub-populations of cells. The regulatory path(s) critical for PlcH induction in the host have not been determined, but based on the current tally of regulators, one might expect there to be some functional redundancy.

The terminator stem-loop downstream of the *cerN* open reading frame is predicted to be a strong Rho-independent terminator with a ΔG of -21.2 kcal/mol (as calculate from ^39^). We were therefore initially surprised that sphingosine induction of *plcHR* occurred via transcriptional activation at the *cerN* promoter. When we tested the *cerN* terminator downstream of a strong heterologous promoter (P_*BAD*_), it showed robust but incomplete repression of readthrough and removal of half of the stem-loop resulted in fully permissive transcription (**Fig. 4A**). Sphingosine does not alter terminator function in *E. coli* or *P. aeruginosa* and we propose that the terminator functions as a static dose regulator controlling the ratio of *cerN* to *plcHR*. The importance of maintaining relative dosing between ceramidase and sphingomyelinase/phospholipase C production is not known.

Induction of *plcH* by GB/GbdR has long been known to be under catabolite repression control^26^, but the direct target of this regulation has not been identified. We show that *plcH* induction by sphingosine is also under catabolite repression control (**Fig. 5**), which likely explains why sphingosine induction of PlcH was not detected in media containing peptone and yeast extract^32^. In contrast, induction of *plcH* by PhoB is not catabolite repressed^26^ and ceramidase expression is sphingosine-inducible even in the presence of rich media^32^. Looking at the sequence in the mRNAs related to *plcH* induction, there are AAnAAnAA motifs that are known to mediate post-transcriptional catabolite repression^40,41^ in the *plcH* 5’ UTR within the PhoB-induced *plcH* transcript and thus not likely functional targets for catabolite repression control. The *cerN* 5’ UTR has an AAnAAnAA motif, as do *gbdR* and *sphR*. Since *cerN* induction by sphingosine is unaffected by rich media, that suggests *cerN* and *sphR* are not direct targets of catabolite repression. Looking at the *cerN-plcH* intergenic region, there are three AAnAAnAA motifs between the *cerN* stop codon and the GbdR-dependent *plcH* transcriptional start site. Catabolite repression thus could impact the *plcHR* side of the conditional operon without directly impacting *cerN* expression, though that prediction has yet to be formally tested. However, it is also important to note that catabolite repression control is often lost in *P. aeruginosa* strains from the CF lung and loss of strong catabolite repression is one of the phenotypes of *lasR* mutants^3,4^.

The *cerN* promoter has been shown to be one of the intergenic loci under pathoadaptive selection in the CF lung^42^. The mutations reported therein are in or just downstream of the SphR binding site, depending on the particular strain. This could suggest selection for modulation of sphingosine-dependent *cerN*, and thus *plcHR*, induction in the context of CF, though the functional consequences of these mutations for *cerN* or *plcH* regulation have not been tested. It is important to note that while CF strains do not lose *plcH* activity, PlcH activity does vary widely, suggesting mechanisms to control the dynamic range of PlcH activity in these strains.

PlcH is not homologous to the best-studied phospholipases C, including those from *Bacillus* and *Clostridium*, but is rather a distinct enzyme in its own PC-PLC/phosphatase class^43^. PlcHs are not unique to *P. aeruginosa*, but it is the species in which most studies of virulence alteration and regulation have been conducted. Within the genus Pseudomonas, *plcH* is carried only by *P. aeruginosa* and two strains of *P. denitrificans*, while strictly environmental and plant pathogenic members lack this hemolytic phospholipase (orthology via ^44^). However, PlcH orthologs are found in a number of bacterial pathogens including *Mycobacterium tuberculosis, Francisella tularensis*, and *Burkholderia pseudomallei*^*43*^. In *B. pseudomallei*, there are two distinct PlcH orthologs encoded in the genome with generally overlapping enzymatic capacity but non-redundant functions in virulence models^45^. Whether regulation of PlcH orthologs in these species is similar to *P. aeruginosa*, including identities of the activating molecules, is mostly unknown, though choline-dependent induction has been shown for *B. pseudomallei*^*45*^.

Regulation of PlcH is multifaceted and there are likely additional unknown pathways to control expression of this important virulence factor. One potential is alternate regulation at the *cerN* promoter, which, in addition to sphingosine, has been shown to be induced in response to PC^32,46^. However, *cerN* is not part of the PhoB regulon^47^, the GbdR regulon^22^, or induced by palmitate^32,46^ or diacylglycerol^32^, pointing to some other moiety of PC as an independent inducing agent for *cerN* and, potentially, *plcH*. Finally, beyond the level of transcription, there may be mechanisms for post-transcriptional, translational, or posttranslational regulation that have not been previously appreciated. While post-transcriptional and translational control have not been reported, PlcH export is mediated by the TAT secretion system^48-50^, the chaperones PlcR1 and PlcR2^51^, and the Xcp/Gsp Type 2 secretion system^49^. There is likely additional regulatory control at one or more of these steps to regulate amount of PlcH exported and alter PlcH localization or activity once exported.

## Materials and Methods

### Strains and growth conditions

*Pseudomonas aeruginosa* PA14 and isogenic mutant strains were maintained on Lysogeny Broth-Lennox formulation (LB) or *Pseudomonas* isolation agar (PIA) plates with 50 μg/mL gentamicin added when appropriate. *Escherichia coli* strains used in this study were maintained on LB plates or liquid LB supplemented with 10 μg/mL or 7 μg/mL gentamicin, respectively. During genetic manipulations, *P. aeruginosa* was selected for, and *E. coli* eliminated, using PIA plates supplemented with 50 μg/mL gentamicin. Prior to transcriptional induction studies, *P. aeruginosa* was grown overnight in morpholinepropanesulfonic acid (MOPS) medium^52^ modified as previously described^53^, and supplemented with 20 mM pyruvate and 5 mM glucose. *E. coli* used for induction studies was grown overnight in MOPS medium with 10% LB (v/v), 5 mM glucose, and 7 μg/mL gentamicin.

### Chemicals and notes on sphingosine stability and solubility

All media and standard chemicals purchased from ThermoFisher or Sigma. Sphingosine was purchased as powder from Avanti Polar Lipids and solubilized in 95% ethanol to make 50 mM stocks. The lot number, stock solution age, and sphingosine deposition into the induction assay all seem to impact the absolute amount of sphingosine-dependent induction, leading to varying levels of induction between experiments shown here. As a lipid with labile function groups, the sphingosine stock solution age appeared to be most important, with ability to induce *plcH* and other sphingosine-responsive genes going down as the stock became older. Evaporation of the ethanol vehicle in multi-well dishes by air-drying vs. a gentle stream of nitrogen gas also showed differential induction, with nitrogen gas drying yielding higher induction levels for similar age stocks.

### Strains, general genetic manipulation, and cloning

All strains and plasmids are listed in **Table 2** and genetic manipulations are described below. All PCRs were conducted using Q5 DNA polymerase (NEB). Primer sequences are listed in the supplemental data (DataSet1, Tab1). Synthetic double-stranded DNA fragments were synthesized by IDT (as gBlocks) and their sequences are listed in the supplemental data (DataSet1, Tab2). All plasmids created for this study were sequenced using Plasmidsaurus (Eugene, Oregon, USA) and select genomes were sequenced using SeqCoast (Portsmouth, New Hampshire, USA). PA14 and PAO1 strains with in-frame chromosomal deletions of *plcHR (PA0843-0844*) and *sphR* (*PA5324*) were generated previously^35^. For plasmid carriage, *P. aeruginosa* and *E. coli* strains were transformed via electroporation and selected for growth on PIA with 50 μg/mL gentamicin and LB with 10 μg/mL gentamicin, respectively.

**Table 2.** Strains and plasmids used in this study.

| Lab Strain ID | Genotype | Plasmid | Source |
| --- | --- | --- | --- |
|  | *E. coli* NEB5 alpha | - | NEB |
| | *E. coli* S17λ*pir* | - | 57 |
| MJ79 | *Pseudomonas aeruginosa* PAO1 wild-type | - | 58 |
| MJ532 | Δ*sphR* in PAO1 | - | 35 |
| MJ143 | Δ*plcHR* in PAO1 | - | 16 |
| MJ984 | *P. aeruginosa* PA14 wild-type (new stock of MJ101) | - | 10 |
| MJ590 | Δ*sphR* in PA14 | - | this study |
| MJ144 | Δ*plcHR* in PA14 | - | this study |
| JR221 | *P. aeruginosa* PA14 wild-type | pMW239 | this study |
| JR223 | Δ*sphR* in PA14 | pMW239 | this study |
| JR186 | *plcH-nLuc-plcR* in PA14 | - | this study |
| JR188 | wild-type revertant from JR186 generation | - | this study |
| JR189 | *plcH-nLuc-plcR* in MJ590 | - | this study |
| JR191 | wild-type revertant from JR189 generation | - | this study |
| JR195 | *PcerN* mutant in PA14 | - | this study |
| JR196 | wild-type revertant from JR195 generation | - | this study |
| JR210 | *PcerN* mutant in PA14 | - | this study |
| JR213 | wild-type revertant from JR210 generation | - | this study |
| JR225 | *cerN-term+rrnB-term, plcH-nLuc-plcR* | - | this study |
| JR227 | wild-type revertant from JR225 generation | - | this study |
| JR243 | PA14 wt (MJ984) | pMW240 | this study |
| JR246 | PA14 wt (MJ984) | pMW241 | this study |

**Plasmids**
| Lab plasmid ID | Description | Source |
| --- | --- | --- |
|  | pUCP22, Pa replicative vector | 59 |
|  | pMQ80, Pa replicative vector | 54 |
|  | pMQ30, allelic exchange | 54 |
| pMW22 | P<sub>plcH</sub>-lacZ<sub>YA</sub> | 23 |
| pMW239 | P<sub>cerN</sub>-*fLuc*-P<sub>plcH</sub>-*nLuc* on pUCP22 | this study |
| pMW240 | P<sub>BAD</sub>-*gfpmut3-cerNterm-nLuc* on pMQ80 | this study |
| pMW241 | P<sub>BAD</sub>-*gfpmut3-cerNterm-mut-nLuc* on pMQ80 | this study |

### Plasmid construction

The pMW22 plasmid was generated in a previous study^23^, while the remaining reporter plasmids were constructed as described below.

The p-P_*cerN*_-*fLuc*-P_*plcH*_-*nLuc* vector was built using by compatible end ligation between a BamH1 cut synthetic fragment PcerN-Fluc-PplcH-nLuc and BamH1 cut and phosphatase-treated pUCP22. Correct cloning and directionality of the insert was determined by EcoR1 digest prior to complete sequencing.

The WT and terminator mutant versions of p-P_*BAD*_*-gfpmut3-cerN-ig-plcH-nLuc* were built using HiFi (NEB) assembly using either synthetic gene fragment cerN-wtTerm-nLuc or cerN-mutTerm-nLuc and HindIII cut pMQ80, which puts the *cerN* terminator and the *cerN-plcH* intergenic-driven *nLuc* downstream of the P_*BAD*_-controlled *gfpmut3* in pMQ80. Diagnostic digest with NdeI was used to assess correct assembly prior to complete sequencing.

### General Allelic Exchange and Chromosomal Alterations

All allelic exchange constructs were generated with upstream and downstream flanking sequences to whichever inserted, deleted, or mutated element was being altered (described in detail below) using the pMQ30 as the non-replicative and counter-selectable vector^54^. After cloning the construct into the pMQ30 backbone, it was transformed into S17 λ*pir* as a conjugative donor. Donor *E. coli* were mixed with their appropriate recipient strain and conjugation allowed to occur in spots on LB plates at 30 °C overnight. Conjugation spots were resuspended and plated on PIA with 50 μg/ml gentamicin to select for single-crossover integrants. After restreaking on PIA with 50 μg/ml gentamicin, single-crossover integrants were moved to LB without selection for 3 h prior to plating on LB with 5% sucrose and without salt at 30 °C overnight. Sucrose resistant colonies were screened for loss of gentamicin resistance prior to PCR screening to determine whether each double-crossover colony was a mutant or WT revertant.

The allelic exchange vector for mutation of the SphR binding site in the *cerN* promoter was built using HiFi assembly (NEB) from two PCR products amplified from PA14 genomic DNA and HindIII and KpnI cut pMQ30. The upstream homology region with the targeted mutation was amplified with primers 2813 and 2815, while the downstream homology region was amplified with primers 2816 and 2814. Correct assembly was assessed by SspI digest prior to complete sequencing. Plasmids matching the predicted sequence were transformed into S17 λ*pir* and allelic exchange conducted as described above. The strains with P_*cerN*_ engineered at the native site were verified by PCR with primers 2817 and 2818, which yielded a 600bp fragment that, in the mutant promoter only, is cleavable by SspI into 448 & 152 bp fragments. This resulted in strain JR195 (P_*cerN*_ mutant in PA14), JR210 (P_*cerN*_ mutant in the plcH-nLuc-plcR background) and their respective wild-type revertants (JR196 and JR213). We sequenced the genome of strain JR195 and its parent strain and the only difference was the anticipated alteration of the *cerN* promoter.

The allelic exchange vector for insertion of *nLuc* as part of the *plcHR* operon was built using HiFi assembly (NEB) from two PCR products generated from PA14 genomic DNA, the gBlock-nLuc-untagged synthetic fragment, and HindIII and KpnI cut pMQ30. The *plcH*-side homology region was amplified with primers 2833 and 2834. The *plcR*-side homology region was amplified with primers 2835 and 2836 for PAO1 and 2837 and 2836 for PA14, to retain strain-specific single nucleotide differences in the *plcH-plcR* intergenic region. Correct assembly was assessed by EcoR1 and NdeI double digest prior to complete sequencing. Plasmids matching the predicted sequence were transformed into S17 λ*pir* and allelic exchange conducted as described above. The strains with *plcH-nLuc-plcR* engineered at their native site were verified by PCR with primers 2896 and 2897 and induction of nLuc expression in response to 2 mM choline (described below), resulting in strains JR186 (PA14 wild-type, *plcH-nLuc-plcR*), JR189 (PA14 Δ*sphR, plcH-nLuc-plcR*), JR189 (PA14 Δ*sphR, plcH-nLuc-plcR*) JR210 (P_*cerN*_ mutation, *plcH-nLuc-plcR*), and JR213 (P_*cerN*_ wild-type revertant, *plcH-nLuc-plcR*). Wild-type revertants (lacking *nLuc*) were saved to use as non-luminescent controls for reporter assays.

The allelic exchange vector for insertion of *rrnB* upstream of the predicted *cerN* transcriptional terminator on the chromosome was built using HiFi assembly (NEB) from two PCR products generated from PA14 genomic DNA, the *rrnB* T1 synthetic fragment, and HindIII and KpnI cut pMQ30. The *cerN*-side homology region was amplified with primers 2846 and 2847. The *plcH*-side homology region was amplified with primers 2848 and 2849. Correct assembly was assessed by BamHI and KpnI double digest prior to complete sequencing. Plasmids matching the predicted sequence were transformed into S17 λ*pir* and allelic exchange conducted as described above. Donor *E. coli* were mixed with recipient PA14 strains JR186 (wild-type, *plcH-nLuc-plcR*) and JR188 (wild-type, *plcHR* revertant). Successful insertion of *rrnB* upstream of the *cerN* transcriptional terminator sequence were then verified by PCR using primers 2853 and 2854, as well as induction of nLuc expression in response to 2 mM choline (described below), generating strains JR225 (*cerN-rrnB-plcH-nLuc-plcR*), JR228 (*cerN-rrnB-plcHR*). Revertant strains, that lack the *rrnB* insertion, were saved to use as controls in reporter assays.

### Induction for reporter and enzymatic assays

Three reporter constructs were used in this study to observe SphR-sphingosine induction of *plcH* transcription: the choromosomal *plcH-nLuc-plcR* reporter, the plasmid-borne pMQ80::P_*BAD*_-*gfpmut3*-ig_*cerNplcH*_-*nLuc* and mutant version reporters, and the plasmid-borne P_*cerN*_-*fLuc*-P_*plcH*_-*nLuc* reporter. *P. aeruginosa* strains were grown overnight in MOPS minimal media with 20 mM sodium pyruvate, 5 mM glucose prior to induction. Cells were collected via centrifugation, washed in MOPS, and resuspended in MOPS with 20 mM pyruvate or in MOPS with 20 mM pyruvate plus 2 mM choline or 100 μM sphingosine (Avanti Polar Lipids). All inductions took place at 37°C and were shaken at 170 rpm for four hours.

### nLuc assay

We observed induction of the chromosomal *plcH-nLuc-plcR* reporter in nanoluciferase reporter assays using the Nano-Glo^®^ Luciferase Assay kit (Promega). The assay was conducted as described by the manufacturer with some modifications reported here. Prior to reading luminescence, induced cells were collected and incubated with 3 mg/mL lysozyme for 30 minutes at 37°C (as the kit does not lyse bacterial cells). Lysed cells were transferred to white (flat clear-bottom) 96-well plates and mixed 1:1 with Nano-Glo^®^ reagent. Luminescence and optical density (OD_600_) were measured using the Synergy 2 H1 Biotek plate reader. The background luminescence of *plcH-plcR* revertant *P. aeruginosa* (no reporter) was subtracted from all values.

Experiments using the plasmid-borne P_*BAD*_-*gfpmut3*-mutig_*cerNplcH*_-*nLuc* reporter were conducted in a similar manner as described above with the addition of measuring GFP fluorescence (485 excitation and 528 nm emission).

### Phospholipase C activity assay

Phospholipase C activity was measured by observing the hydrolysis of the synthetic substrate ρ-nitrophenyl-phosphorylcholine (NPPC) based upon the Kurioka and Matsuda method ^55^, modified as we have described previously ^23^, but using a final concentration of 10 mM NPPC. Briefly, *P. aeruginosa* was grown overnight in MOPS minimal media with 20 mM sodium pyruvate, gentamicin, and 0.05% L-arabinose; with or without 1 mM choline and with or without added NaCl. NPPC hydrolysis was measured by monitoring the absorbance at 410 nm over time. Phospholipase C activity was reported in micromoles of ρ-nitrophenol generated per minute of reaction per optical density (OD_600_). ρ-nitrophenol concentration was calculated using the extinction coefficient of 17700 M^-1^ cm^-1 56^.

### RT-PCR for operon determination

RNA was isolated as described above from PA14 induced in MOPS pyruvate, MOPS pyruvate with 100 μM sphingosine, and MOPS pyruvate with 2 mM choline. cDNA was generated using the Induro RT reverse transcriptase system (NEB). PCRs were conducted using Q5 DNA polymerase (NEB) for *cerN* (primers 2850 & 2851), *plcH* (primers 2858 & 2859), and the *cerN-plcH* intergenic region (primers 2866 & 2867) and products separated by agarose gel electrophoresis. PA14 genomic DNA was the positive amplification control, while the negative amplification control was no added template. Control for DNA contamination of the RNA was done by leaving out the reverse transcriptase from the cDNA creation reaction.

### Data acquisition, data analysis, data visualization, and statistics

All absorbance, luminescence, and fluorescence measures were done using either BioTek Synergy2 or BioTek H1 multimode plate readers and data acquired using Gen5 (BioTek). Data were exported to Microsoft Excel and all normalization and fold-change calculations were done in Excel. The individual data values used in each figure are available in Supplementary material (DataSet 2). For visualization and statistical analysis, data were moved to GraphPad Prism. Specific statistical tests and their comparator groups are noted in the figure legends. We have opted to show all collected data points using color coding to separate the different experiments however, and importantly, statistical testing was done only using the means from each experiment. Diagrams of chromosomal loci and plasmid constructs were generated in BioRender and used here under the academic licensing terms.

## Acknowledgements

The Vermont Integrative Genomics Resource Massively Parallel Sequencing Facility was supported by the University of Vermont Cancer Center, Lake Champlain Cancer Research Organization, UVM College of Agriculture and Life Sciences, and the UVM Larner College of Medicine.

## Notes

### Competing Interest Statement

The authors have declared no competing interest.

